# Head motion during fMRI tasks is reduced in children and adults if participants take breaks

**DOI:** 10.1101/816116

**Authors:** Tobias W. Meissner, Jon Walbrin, Marisa Nordt, Kami Koldewyn, Sarah Weigelt

## Abstract

Head motion remains a challenging confound in functional magnetic resonance imaging (fMRI) studies of both children and adults. Most pediatric neuroimaging labs have developed experience-based, child-friendly standards concerning e.g. the maximum length of a session or the time between mock scanner training and actual scanning. However, it is unclear which factors of child-friendly neuroimaging approaches are effective in reducing head motion. Here, we investigate three main factors including (i) time lag of mock scanner training to the actual scan, (ii) prior scan time, and (iii) task engagement in a dataset of 77 children (aged 6-13) and 64 adults (aged 18-35) using a multilevel modeling approach. In children, distributing fMRI data acquisition across multiple same-day sessions reduces head motion. In adults, motion is reduced after inside-scanner breaks. Despite these positive effects of splitting up data acquisition, motion increases over the course of a study as well as over the course of a run in both children and adults. Our results suggest that splitting up fMRI data acquisition is an effective tool to reduce head motion in general. At the same time, different ways of splitting up data acquisition benefit children and adults.

**Highlights:**

- In children, fMRI data acquisition split into multiple sessions reduces head motion
- In adults, fMRI data acquisition split by inside-scanner breaks reduces head motion
- In both children and adults, motion increases over the duration of a study
- In both children and adults, motion increases over the duration of a run

## 1 Introduction

“Please remember: Relax and try not to move. Here we go.” In functional magnetic resonance imaging (fMRI) experiments, this is often the last thing the researcher says before starting the scanner and the experiment. What is left to do is hoping for good quality data. As MRI is very susceptible to head motion during the scan (Friston et al., 1996), hoping for good quality data is often equivalent to hoping for data with low head motion. The importance of reducing head motion during data acquisition has been impressively documented in several studies showing that head motion can lead to misleading results (in some cases, even after retrospective motion correction; Power et al., 2012; Satterthwaite et al., 2012; van Dijk et al., 2012).

In pediatric neuroimaging studies, the general problem of head motion is especially challenging. Despite children’s high motivation to lie still, in most studies children still move more than adults. This poses a problem for data quality in pediatric neuroimaging especially for group comparisons between children and adults. Even with retrospective head motion correction (which is limited in its ability to correct for motion Field et al., 2000; Freire and Mangin, 2001; Friston et al., 1996), group differences in head motion often persist. This usually leads to the exclusion of motion-affected runs or participants from data analysis (e.g. Meissner et al., 2019; Nordt et al., 2018; Walbrin et al., 2020), hence requiring additional time and research funds to achieve an adequately powered study design.

To minimize motion during data acquisition, various solutions have been developed. For structural MRI scans in clinical settings, sedation is often used. However, this is not an option for fMRI studies in research settings due to the need for attentive participants as well as the unacceptable risk of health-related side effects (Bie et al., 2010). One set of solutions for fMRI studies aims at restraining head motion physically. Restrictive approaches including bite bars (Menon et al., 1997) or thermoplastic face masks (Green et al., 1994), are considered effective for short scan durations. However, they reduce comfort and are therefore seldomly accepted in pediatric neuroimaging. In contrast to the desired outcome, the discomfort of these methods can also lead to more fidgeting and wiggling in search of comfort (Zeffiro, 1996), especially for longer scans. Approaches that use individually 3D-printed styrofoam head molds are effective and said to be comfortable (Power et al., 2019), but require additional time and research funds for each participant. A less resource-intensive, yet effective and accepted solution, is to provide tactile feedback about participant’s head motion by applying a tape or ribbon across the head coil that touches the participant’s forehead (Krause et al., 2019).

Aside from physical constraints, pediatric neuroimaging groups often adapt the study procedure to reduce children’s motion during data acquisition. The most prominent tool is to precede the actual MRI session(s) with a scanner training session (Raschle et al., 2009; Slifer et al., 1993). Scanner training sessions are usually performed in a mock scanner, a custom-built or purchased facsimile of a real MRI, which lacks the technical capability to acquire actual data, but can play scanner sounds and has a similar setup to the real scanner (e.g. head coil-mounted mirror, response buttons, etc.). During these sessions, behavioral training is used to teach lying still in the scanner bore and responding to the task. There is a general consensus that a scanner training session is beneficial for fMRI data acquisition as it reduces children’s anxiety (Durston et al., 2009; Raschle et al., 2009; Rosenberg et al., 1997). While a reducing effect on head motion is also assumed, the current literature has not been able to show this (due to a lack of control groups that did not receive scanner training; but see Barnea-Goraly et al., 2014; Bie et al., 2010; Epstein et al., 2007 for success rates and reduced motion after scanner training).

Similarly, many pediatric neuroimaging groups have established experience-based guidelines for child-friendly study designs in terms of the scanning procedure. These usually include keeping runs short, interspersing anatomical scans as task-free inside-scanner breaks while presenting entertaining video clips, and limiting the total scanning session time. Moreover, based on findings in adults (Huijbers et al., 2017), tasks are designed in an engaging and interactive way rather than just requiring passive perception of stimuli. In consequence, a small body of literature has developed that recommends scanning procedures for pediatric neuroimaging (e.g. Greene et al., 2016; Habibi et al., 2015; Raschle et al., 2009). However, as for scanner training, the contribution of presumably child-friendly adaptations in scanning procedures on minimizing head motion has not been investigated so far.

The present study set out to identify which fMRI study procedures contribute to reduced head motion in fMRI studies. Our aim was to generate data-driven suggestions to optimize study procedures for children and adults separately. To this end we utilized head motion estimates derived from a standard motion correction pipeline applied to 77 children and 64 adults from three fMRI studies at two sites, and notes that yielded demographic data, the date of scanner training, and the sequence of data acquisition. Using separate multilevel linear models for children and adults, we investigated the effect of splitting up data acquisition into several days or sessions, the effect of interspersing functional data acquisition with structural runs and video clip breaks, the effect of time between scanner training and the actual scan, and the effect of task engagement.

## 2 Methods

### 2.1 Definition of session, run, and functional segment

In this article, we make important distinctions between “session”, “run”, and “functional segment”. Session corresponds to a continuous period of time spent inside the scanner. A session begins upon entering the scanner and ends as a participant leaves the scanner. For example, if a participant enters the scanner, takes a break outside the scanner and re-enters the scanner, this would constitute two sessions. Run corresponds to a continuous image acquisition sequence. For example, a participant could complete an experiment with four fMRI runs within a single session. Functional segment corresponds to a period of consecutive acquisition of functional runs inside the scanner. For example, if a scan procedure involves an anatomical T1-scan, three fMRI runs, two diffusion-weighed imaging (DWI) sequences, and finally four more fMRI runs, this participant has completed two functional segments.

### 2.2 Participants

Our study included data from two neuroimaging centers (see 2.3, Neuroimaging) and three developmental cognitive neuroimaging studies, a total 680 runs from 77 children and 624 runs from 64 adults. Some of the data has been used to answer questions concerning the neurocognitive visual and social development in children and adults previously (Meissner et al., 2019; Nordt et al., 2018; Walbrin et al., 2020). The final analyzed data set was reduced to 626 runs from 77 children and 470 runs from 54 adults due to several exclusion criteria (see 2.5, Data exclusion). In children, the number of runs per participant ranged from 1 – 14 (*M* = 8.13, *SD* = 3.21), in adults, the number of runs per participant ranged from 4 – 14 (*M* = 8.70, *SD* = 3.33). Children’s age ranged from 6.78 – 13.01 years (*M* = 9.52, *SD* = 1.68), adult’s age ranged from 18.41 – 35.02 years (*M* = 21.76, *SD* = 2.67). All participants were healthy, had normal or corrected-to-normal vision, and had been born at term. No participant reported past or current neurological or psychiatric conditions, or had structural brain abnormalities. All participants as well as children’s parents gave informed and written consent to participate voluntarily.

### 2.3 Neuroimaging

fMRI took place at the Neuroimaging Centre of the Research Department of Neuroscience at Ruhr University Bochum and the Bangor Imaging Centre at Bangor University. At Bochum, a total of 338 runs in 50 children and 346 runs in 40 adults were acquired—at Bangor, a total of 188 runs in 27 children and 123 runs in 14 adults were acquired. At both sites, 3.0T Achieva scanners (Philips, Amsterdam, The Netherlands) and 32-channel head coils were used (Supplementary Figure S1). For all fMRI, we used blood oxygen level dependent (BOLD) sensitive T2*-weighted sequences. Across sites and experiments, the following fMRI acquisition parameters were constant: FOV = 240 mm × 240 mm, matrix size = 80 × 80, voxel size = 3 mm × 3 mm × 3 mm, TR = 2000 ms, TE = 30 ms. However, slice gap (Bangor: 0 mm, Bochum: 0.4 mm), number of slices (Bangor: 32, Bochum: 33), slice scan order (Bangor: ascending, Bochum: ascending interleaved), and flip angle (Bangor: 83°, Bochum: 90°) differed between sites. Visual stimuli were presented via a VisuaStim Digital goggle system (FOV: 30° × 24°, 800 × 600 pixel, Resonance Technology Inc., CA, USA) in Bochum and via a MR-safe monitor and mirror system (FOV: 25.92° × 16.90°, 1920 × 1200 pixel; Cambridge Research Systems, UK) in Bangor. Due to the visual isolation inherent with the goggle system, a researcher was present next to the scanner bore opening at all times for immediate verbal contact and motion feedback in Bochum, but not in Bangor.

For each study, a certain number of functional runs and structural scans were planned. Details on the experiments that were done during the functional runs as well as the number of participants and runs for each of the tasks are given in the Supplementary Text S1 and Supplementary Table S1. In adults, experimenters followed this protocol, checking in with participants after the completion of each experiment (i.e. after multiple runs) to explain what would happen next and only stopped or diverted from protocol if participants actively reported feeling unwell. In contrast, in children, we checked in with participants after each run to actively inquire about their well-being. Moreover, nearing the end of a session, we also actively asked if they still felt good and ready to do another run. This was done in order to give children the opportunity to express any signs of discomfort, which to our experience children do not necessarily utter spontaneously but sometimes only after encouraging them to be open about it or giving them an explicit opportunity. Thus, we dynamically decided when to break up acquisition into sessions or days, when to intersperse tasks with a structural scan accompanied by an entertaining video, or when to end the study.

### 2.4 Scanner training

All children underwent a scanner training in order to familiarize them with the scanner environment and achieve high-quality scans with as little motion as possible (Supplementary Figure S1). At both scanner sites, pre-recorded audio of MRI acquisition sequences was played during scanner training to simulate the real scanner environment and visual stimulation was achieved through a mirror system targeted at a monitor outside of the mock scanner bore.

In other aspects, training sessions differed between sites. In Bangor, scanner training was conducted right before scanning and entailed lying inside a mock scanner with a motion sensitive electrode placed on the forehead to measure movement across three translation and three rotation axes (MoTrak Head Motion Tracking System; Psychology Software Tools, 2017). Children received visual motion feedback via an on-screen cursor that was controlled by children’s head movements. Children were instructed to lie still and keep the cursor in the middle of a target circle, whose diameter allowed for 3 mm head translations in any direction. Once children were able to keep the cursor within this target region for a timed period of 30 seconds, children watched a short animated video. When movement exceeded 3 mm translation, video playback was paused, providing immediate feedback that they had moved too much. Once children were able to watch the video for a period of 2 minutes without a video pause, scanner training was completed.

In Bochum, scanner training was conducted between one to ten days before the MRI study for the majority of participants (n = 48). A minority was trained on the same day as the first scan (n = 1) or between 12 to 35 days before the first scan (n = 7). Training began with sitting on the extended mock scanner bed, watching a short animated video that introduced a cover-story explaining the tasks to be performed inside the scanner, and practicing these tasks (button-press). This was followed by explaining and demonstrating the procedure of entering an MRI scanner with a large puppet. Next, children entered the mock scanner bore, watched a short animated video and performed several practice trials of the tasks presented on a screen that was visible via a mirror system. The researchers gave feedback with respect to the level of movement and task performance throughout the scanner training session. It was established that a researcher would gently touch the children’s shin in case of excessive motion as a means of motion feedback. Verbal feedback was gradually decreased throughout the session. As head motion during the mock scanner session in Bochum was not quantified, feedback was based on the experimenter’s observation of children’s head motion. Once the researcher decided that motion levels were acceptable and the tasks were understood, scanner training was completed.

### 2.5 Data exclusion

Our initial dataset encompassed all completed fMRI runs of two developmental cognitive neuroscience studies conducted in Bochum and one in Bangor. Aborted runs (e.g. due to technical errors) were not included, as they were not retrieved from the MRI-controlling computer and not reliably recorded in handwritten notes. To ensure that our results are valid and interpretable, we excluded data that would have biased our analyses (Figure 1).

**Figure 1:**
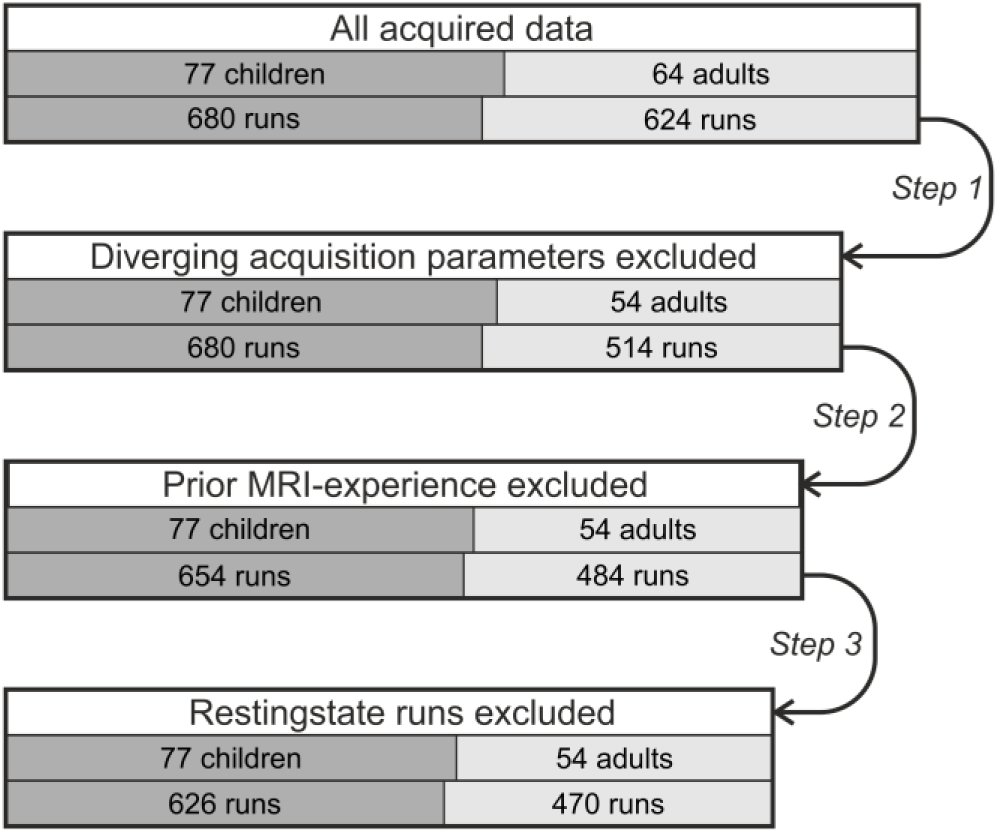
Data exclusion steps and resulting sample sizes for children (left, dark gray) and adults (right, light gray). Main box widths are in relation to the original total number of runs. Box segmentation mirrors the relative distribution of adult vs. child participants or runs, respectively.

In a first step, we excluded ten participants (110 runs) from Bangor that were recorded with different acquisition parameters, i.e. with a slice thickness of 3.5 mm instead of 3.0 mm. This difference in voxel size along the z-axis would affect three out of the six motion estimation parameters (rotation around the y-axis, rotation around the x-axis, and translation along the z-axis) during motion correction, leading to systematically larger motion estimates. In a second step, we excluded 56 runs that did not reflect first-time fMRI experience. Four participants had participated in both Bochum fMRI studies that are included in our analysis. Participation in both studies was separated by 2-18 months. To control for possible confounding effects of prior scanning experience on motion, we excluded all runs of their second fMRI study participation. Implementing this control step ensured that we report on the rather typical case—at least for children—of a first-time MRI study participant. Note that we could not rule out the possibility that participants took part in an MRI study of a different lab before, although this is highly unlikely for children and most adults at our neuroimaging centers. In a last step, we excluded 42 resting-state runs. Resting-state runs were only acquired in one of the three studies, only once per participant, and in the majority of cases the run was acquired at the end of a session. Moreover, resting-state runs were the only runs in which participants had their eyes closed during acquisition. This unique set of circumstances makes resting-state runs very likely to bias our results, as there is not enough data to efficiently model this set of circumstances, e.g. as an independent variable.

### 2.6 Head motion estimation

To estimate head motion during fMRI scans, we used the neuroimaging software package BrainVoyager (Version 20.2 for 64bit Windows, RRID: SCR_013057). First, fMRI run series in DICOM format were converted to the proprietary STC and FMR format using BrainVoyager scripts. Then, we applied BrainVoyager fMRI data preprocessing tools in their default settings. That is, for slice scan time correction, we used cubic spline interpolation. For 3D motion correction, we used a trilinear detection and sinc interpolation approach with a maximum of 100 iterations to fit a volume to the reference volume. Resulting motion log files contained six timeseries representing the estimated volume-wise instantaneous translation and rotation for axes x, y, and z in reference to the first volume of the respective experiment.

For each of the six motion parameters, we calculated a timeseries of volume-to-volume motion, i.e. the difference between each volume’s motion parameter and the previous volume’s motion parameter. Rotational motion was converted from degree to millimeter by calculating displacement on the surface of a sphere of 50 mm radius (approximating the distance from the center of the head to the cortex; Power et al., 2012). The framewise displacement of the head position (FD) was calculated as the sum of the absolute volume-to-volume motion values (Power et al., 2012). FD was shown to have a strong association with motion-induced artifacts (Ciric et al., 2017). For each run, we calculated the mean FD.

Moreover, for discrete one-minute-intervals (i.e. 30 volumes) within runs, we calculated the percentage of high-motion volumes with FD above the threshold of 0.3 mm (volume-to-volume). This threshold for high-motion volumes was determined by extensive exploration of our data before any statistical analysis, and aimed at capturing motion “spikes” without impacting the motion “floor” volumes as done previously (Power et al., 2019; Power et al., 2012) and resulted in plausible ratios of high-motion volumes for children (*M* = 12.80 %, *SD* = 20.89 %) and adults (*M* = 4.48 %, *SD* = 12.10 %). For a visualization of the high-motion volume threshold, see Figure 2. The two head motion measures—mean FD per run and frequency of high-motion volumes per minute of a run—were investigated in separate analyses.

**Figure 2:**
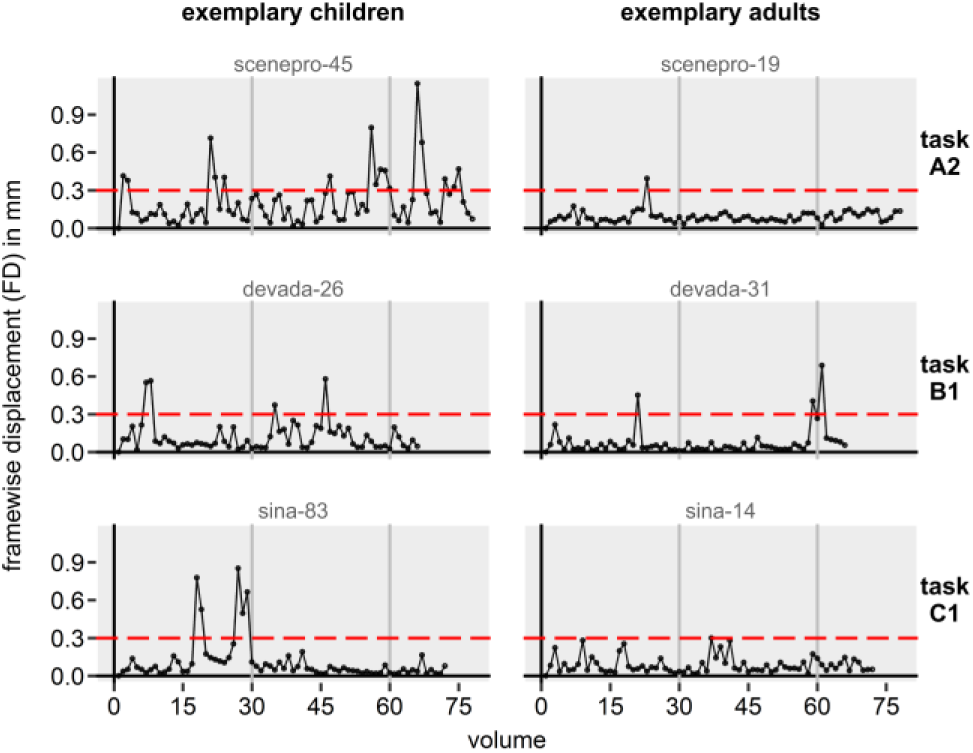
Framewise displacement (FD) of the head in mm in three example children and adults (gray writing = participant IDs). For each participants, one representative run was chosen to show the high-motion volume threshold (red horizontal dashed line). Gray vertical lines indicate the one-minute-intervals, for which the percentage of high-motion-volumes was calculated.

### 2.7 Predictor variables

Our study did not aim to examine age group differences between children and adults, but to provide guidelines for practitioners and researchers who conduct pediatric neuroimaging examinations and/or experiments that might also include adult control groups. Thus, we report separate analyses for children and adults. This approach enabled us to include two more predictor variables (PVs) for children, as adults did not perform scanner training and we did not expect age effects in our range of 18-35-year-old adults. For each age group, we assessed two head motion measures with up to eight possible predictors.

#### 2.7.1 Mean FD

In our first analysis, we asked which factors would influence the mean motion during a run. Seven PVs were evaluated:

- PVs 1-4) *Prior scan time* encoded the time in minutes that a participant had been in the scanner already. That is, the summed scan time of all functional as well as structural runs that had been administered 1) since the beginning of the *functional segment*, 2) since the beginning of the *session*, 3) since the beginning of the *day*, or 4) since the beginning of the *study*. For definitions of *session* and *functional segment*, please refer to section 2.1, Definition of session, run, and functional segment. Note that *prior scan time since the beginning of the day* was only analyzed in children, because while children were scanned on either one (n = 45), two (n = 31) or three days (n = 1), only 2 out of 54 adults were scanned on multiple (here: two) days. Therefore, our adult data did not have the required distribution to allow any inferences on breaking up fMRI data acquisition between days in adults. For example, imagine the complex case of a participant that comes in for a second day of fMRI scanning (after 60 min. of scanning on the first day), takes a bathroom break after the completion of the day’s first fMRI experiment (after 15 min.), re-enters the scanner, completes the second fMRI experiment (15 min.), and is now scheduled for an inside-scanner break, during which a diffusion-weighed imaging sequence is being acquired while a video clip is presented (5 min.). The next fMRI run that is acquired would have different values for each of the *prior scan time variables*, e.g. a) *prior functional segment scan time* = 0 min., b) *prior session scan time* = 20 min., c) *prior day scan time* = 35 min., and d) *prior study scan time* = 95 min. In general, we hypothesized that participants’ motion would increase with time spent in the scanner, but could be “reset” to lower motion by breaks. Thus, we tested the effect of four different *prior scan time* variables to determine which—if any—way of breaking up data acquisition reduces subsequent motion. Figure 3 visualizes the possible main effects of *prior scan time* variables. To determine the *prior scan time* variables values, we evaluated structured handwritten MRI session notes that stated the sequence of acquired functional and structural runs and breaks between sessions and days for each participant. Then, the duration of each run was retrieved from its DICOM file header. This DICOM header duration always exceeded the product of repetition time and number of volumes, i.e. it included scanner- and sequence-specific preparation time. We did not include potential breaks between runs that occur due to normal operation time that is required to start a new paradigm on the stimulation computer or to start the scanner, how-do-you-feel-inquiries, or minor technical issues, because we did not log these events. Further, we did not include aborted runs, because we did not save this data nor took reliable notes, rendering a determination of the duration of these scans impossible.
- PV 5) *Age* was defined in years at the first day of scanning and was only investigated for children. This variable was included to explore possible interactions with the other PVs, not for main effects of age, as main effects of age are well established in the field and would only lead to an unnecessarily complex model with less power to detect effects of interest.
- PV 6) *Scanner training date* was recorded as the number of days that passed since the scanner training for children only. We hypothesized that a greater time interval between scanner training and actual MRI scan would result in higher motion.
- PV 7) *Task engagement* encoded how much active engagement a given fMRI run required. We distinguished if participants just had to passively watch the display, or if participants had to perform a task that included button-pressing. We expected that an active task engagement would result in reduced motion due to enhanced attention and less awareness of a possibly uncomfortable situation, itching, or other distractions.

**Figure 3:**
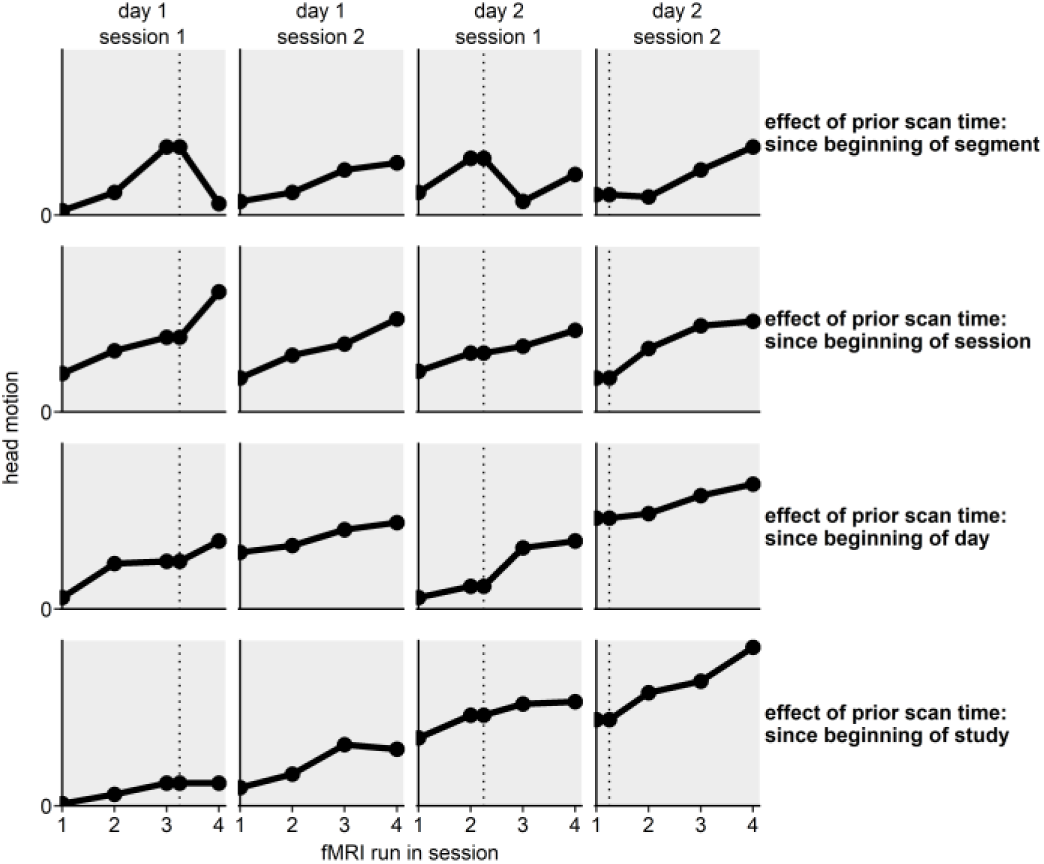
Visualization of the effect of prior scan time with hypothetical data. Each row shows the effect of a different kind of prior scan time. Each column shows a single session, consisting of four fMRI runs. The dotted vertical line and points on the vertical line denote inside scanner breaks during which structural scans were acquired and video clips were presented.

In addition to these main effects, we investigated the possible interactions between the *prior scan time* variables and the other variables. Other two-way or three-way interactions were not investigated to avoid overly complex and highly computation-intensive models and as the detection of high-order effects would require a more powerful study design (Simonsohn, 2014).

#### 2.7.2 High-motion volumes

In our second analysis we asked which variables would influence head motion over the course of a run. As decisions on retaining or discarding runs for further analysis are often informed by high-motion volumes within a run rather than a run’s mean motion, instead of mean FD, we investigated the percentage of high-motion volumes for each minute of a run (i.e. FD_volume_ > 0.3 mm; see 2.6, Head motion estimation). Consequently, eight PVs were evaluated:

- PVs 1-7) All previous PVs, i.e. *prior functional segment scan time, prior session scan time, prior day scan time, prior study scan time, age, scanner training date, task engagement*
- PV 8) *Minute of run* coded discrete one-minute-intervals within runs, allowing us to investigate the course of high-volume motion occurrence across the course of a run.

In addition to main effects, we investigated the possible interactions between *minute of run* and the other variables. Again, we did not investigate other two- or three-way interactions (see 2.7.1, Mean FD)

### 2.8 Statistical model

#### 2.8.1 Hierarchical data structure

We analyzed our data using multilevel linear models (MLMs) due to the hierarchical four- or five-level grouping structure inherent in our data (cf. Engelhardt et al., 2017; Figure 4): Motion estimations for mean FD per run are nested within sessions (level 4), sessions are nested within days (level 3), days are nested within participants (level 2), and participants are nested within studies (level 1). Motion estimations for percentage of high-motion volumes per minute of a run are nested within runs, establishing a fifth level. This hierarchical data structure is likely to introduce dependency of observations within a grouping variable. For example, motion from two randomly selected runs from the same participant is likely to be much more similar than motion from different participants. In technical terms, the residuals within a grouping variable are likely to be correlated. Thus, the assumption of independent residuals, crucial for parametric tests, is violated. While dependent observations for one level, e.g. within participants, could be handled easily by a conventional repeated-measures ANOVA, this case is a complex nested multilevel dependency structure that cannot be adequately and validly handled by a repeated-measures ANOVA—but can be by an MLM.

**Figure 4:**
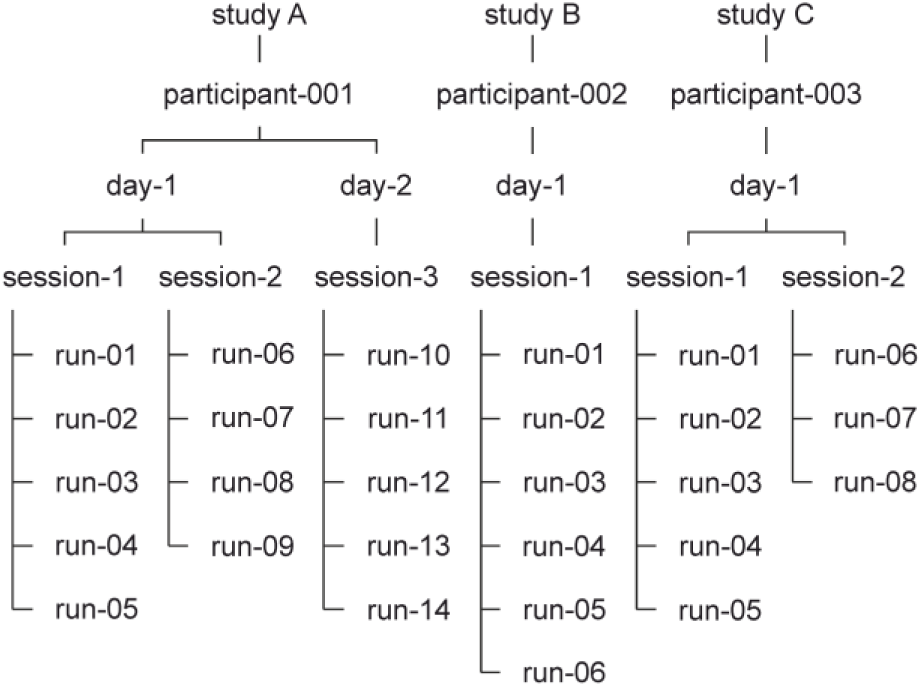
Hierarchical data structure with four levels. For visualization purposes, only one fictional subject is displayed for each study.

#### 2.8.2 MLM creation step 1: Identifying relevant grouping structures

MLMs were created in a data-driven process. First, we assessed the possibility that the grouping variables would introduce dependencies in the data—and thus confirm the need for an MLM. To this end, we calculated the intraclass correlation (ICC, i.e. the proportion of the total variance that is explained by the respective grouping factor) for each grouping level of the model and liberally incorporated all grouping levels into our model that would explain at least 1% of the total variance (i.e. ICC ≥ 0.01, Supplementary Text S2). Next, we used the chi-square likelihood ratio test and Akaike’s information criterion (AIC) to test if the model that incorporated the grouping structure actually had a better fit to our data than the model without the grouping structure and included or ignored the grouping structure in all subsequent models accordingly (Supplementary Text S3).

ICC analysis for mean FD over the course of a study indicated that in both children and adults, the grouping factors *participant* (children: 0.487, adults: 0.376) and *session* (children: 0.229, adults: 0.021) explained a substantial amount of variance in the data, while the grouping factors *study* and *day* did not (both age groups < 0.001). In both children and adults, models that allowed random intercepts for *participant* and *session* fit the data significantly better than a model with fixed intercepts (children: Supplementary Table S4, adults: Supplementary Table S5). Thus, the MLMs were built with random intercepts for *participant* and *session*.

ICCs for the percentage of high-motion volumes across the course of a run in both children and adults revealed substantial dependency within the grouping levels *participant* (children: 0.136, adults: 0.292), *session* (children: 0.275, adults: 0.037), and *run* (children: 0.143, adults: 0.309), but not within the grouping levels *study* (both age groups: < 0.001) and *day* (children: < 0.001, adults: 0.004). A random intercept model for *participant*, *session*, and *run* fit our data better that a fixed intercept model (children: Supplementary Table S6, adults: Supplementary Table S7). Consequentially, the MLMs were built with random intercepts for *participant*, *session* and *run*.

#### 2.8.3 MLM creation step 2: Stepwise inclusion of relevant predictors

Second, we introduced the PVs, i.e. main effects and selected interactions (fixed effects), into the model stepwise, to test if they improved the model fit significantly. The order of introduction for mean FD and high-motion volumes MLMs is listed in Supplementary Table S2 and Supplementary Table S3, respectively. If a model including a fixed predictor led to a better model fit than the previous model without it, the predictor was included in all subsequent models, otherwise it was left out of subsequent models.

For children’s mean FD MLM main effects of *prior functional segment time*, *prior session time*, *prior day time*, and *scanner training date*, as well as the interaction effects of *age* × *prior session time* and *age* × *prior study time* were included in the model (Supplementary Table S4). For children’s high-motion volumes MLM, main effects of *minute of run*, *prior segment time*, *prior session time*, *prior day time*, and *scanner training date* were added to the model (Supplementary Table S6). For adult’s mean FD MLM, main effects of *prior functional segment time*, *prior study time*, and a *prior functional segment time* × *task engagement* interaction were added to the model (Supplementary Table S5). For adult’s high-motion volumes MLM main effects of *minute of run*, *prior segment time*, and *prior session time*, were added to the model (Supplementary Table S7).

#### 2.8.4 MLM creation step 3: Assessing the need for participant-specific predictor effects

Third, we tested if the model fit improved further if we let the effect of each included main effect predictor vary over participants (random slopes) and added significant random effects to the model accordingly. Predictors *scanner training date* and *age* were not considered as they are constant for each participant’s run. Further, we did not fit random slopes for interaction terms as this prevented the MLMs to converge—possibly because our data did not have enough power to fit a model of such high level of complexity.

Adding random slopes across participants for all main effects further improved model fit in all MLMs and thus was incorporated in the final models (Supplementary Table S4, S5, S6, S7).

### 2.9 Software

Data handling and statistical data analysis was performed using R (version 3.6.0, RRID: SCR_001905, R Core Team, 2015) in RStudio (version 1.2.1335; RRID: SCR_000432). For MLMs, we used the nlme package (version 3.1-140, RRID:SCR_015655, Pinheiro et al., 2019). Figures were created using the ggplot2 package (version 3.2.0, RRID:SCR_014601, Wickham, 2016). Regression lines and confidence interval bands for fixed effects visualizations, i.e. predictor effect plots, were created using the effects package (version 4.1-1, Fox and Weisberg, 2018). Marginal R^2^ (for fixed effects) and conditional R^2^ (for fixed effects and random effects combined) goodness-of-fit was estimated using the r.squaredGLMM function of the MuMIn package (version 1.43.6, Barton, 2019), which is based on methods by Nakagawa and Schielzeth, 2013, Johnson, 2014, and Nakagawa et al., 2017.

## 3 Results

We evaluated which variables would predict head motion during fMRI in children and adults. We assessed two measures of head motion in separate analyses: First, we investigated the course of head motion across an fMRI study in terms of mean framewise displacement per run. Then, we investigated the course of head motion over the course of a run in terms of the frequency of high-motion volumes per minute. Each analysis was performed separately for children and adults. Regression coefficients for all main and interaction effects included in the MLMs were tested against zero using one-sample t-tests with a significance threshold of α = 0.05.

### 3.1 Splitting up data acquisition reduces motion

#### 3.1.1 Children

Evaluation of fixed effects predictors in the final mean FD MLM revealed that *prior session scan time* and a *prior study scan time* × *age* interaction significantly predicted FD in children (Figure 5, Table 1). Children showed greater motion with ongoing session length (Figure 5, left). Note that the total number of sessions in children ranged from 1 – 4 (16 children did only one session, 55 children did two sessions, four children did three sessions, and two children did four sessions). In addition, motion increased with study length in older children, but not as much in younger children (Figure 5, right). As our data in children shows one obvious outlier, a participant whose mean FD is around 2 mm or all of her/his five runs, we tested if this case would influence the observed effects. However, a reanalysis without this participant only slightly increased the effect of prior session scan time and did not change the *age* × *prior study scan time* interaction notably. Thus, here, we report the full data set.

**Figure 5:**
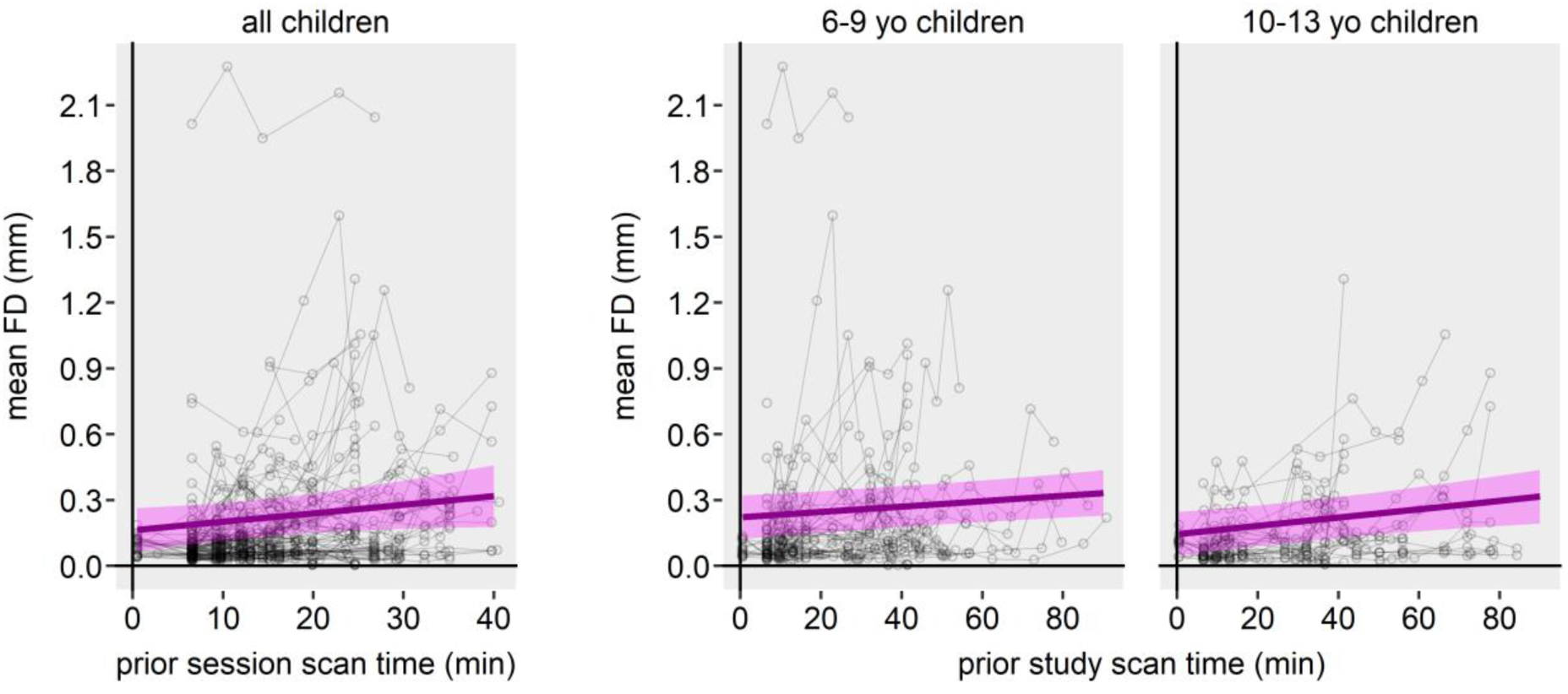
Predictor effect plots for children’s head motion across the course of a study.Thick magenta lines are fitted linear regression lines for the fixed effects predictors denoted on the x-axes. Shaded magenta areas show the 95% confidence band for the regression lines. Each gray circle represents one run. Left: Head motion across the course of a session. Each circle-connecting line represents a participant‘s session. Participants that were scanned in multiple sessions are represented by multiple lines. Right: Head motion across the course of a study for 6-9-year-old and 10-13-year old children. Age bins were chosen arbitrarily to visualize the age × prior study scan time interaction. Each circle-connecting line represents one participant.

**Table 1.** Fixed effects parameter estimates of final model for children’s mean motion across the course of a study.

| parameter | $\beta$ | Lower 95% CI | Upper 95% CI | SE | df | t | p |
| --- | --- | --- | --- | --- | --- | --- | --- |
| Intercept | 0.04895 | 0.01800 | 0.07990 | 0.01584 | 509 | 3.090 | 0.002 |
| Prior functional segment scan time | 0.00147 | -0.00170 | 0.00464 | 0.00162 | 509 | 0.904 | 0.366 |
| Prior session scan time | 0.01564 | 0.00307 | 0.02821 | 0.00644 | 509 | 2.430 | 0.015 |
| Prior day scan time | 0.00212 | -0.00043 | 0.00467 | 0.00130 | 509 | 1.627 | 0.104 |
| Scanner training date | 0.00560 | -0.01790 | 0.02909 | 0.01203 | 509 | 0.465 | 0.642 |
| Age × prior session scan time | -0.00121 | -0.00245 | 0.00003 | 0.00063 | 509 | -1.905 | 0.057 |
| Age × prior study scan time | 0.00016 | 0.00005 | 0.00028 | 0.00006 | 509 | 2.721 | 0.007 |

Note: CI = confidence interval, SE = standard error, df = degrees of freedom

#### 3.1.2 Adults

For adults, evaluation of fixed effects predictors in the final model revealed that *prior functional segment scan time* and *prior study scan time* significantly predicted motion (Figure 6, Table 2). Adults’ motion increased with ongoing *study scan time* (note that in 52/54 adults *study scan time* is equal to *day scan time*). Moreover, the longer functional scans are acquired without an inside-scanner break (e.g. through a structural scan with relaxing video), the more adults moved. Originally, we also found that a *task engagement* × *prior functional segment scan time* interaction significantly predicted motion, i.e. for passive tasks, adults showed a steeper increase in motion than for active tasks (Figure 6, middle, thick black dashed line). However, as it seemed possible that this *task engagement* × *prior functional segment scan time* interaction was driven by an extreme increase in motion at the end of a functional segment in passive tasks in a single participant only (Figure 6, thin black dashed lines and squares), we reanalyzed the MLM without data from this participant. While all other effects remained stable (Figure 6, see overlapping green solid and black dashed lines), the *task engagement* × *prior functional segment scan time* interaction was rendered insignificant by the exclusion of the participant (Figure 6, see black dashed line outside of green confidence band, Supplementary Table S8).

**Figure 6:**
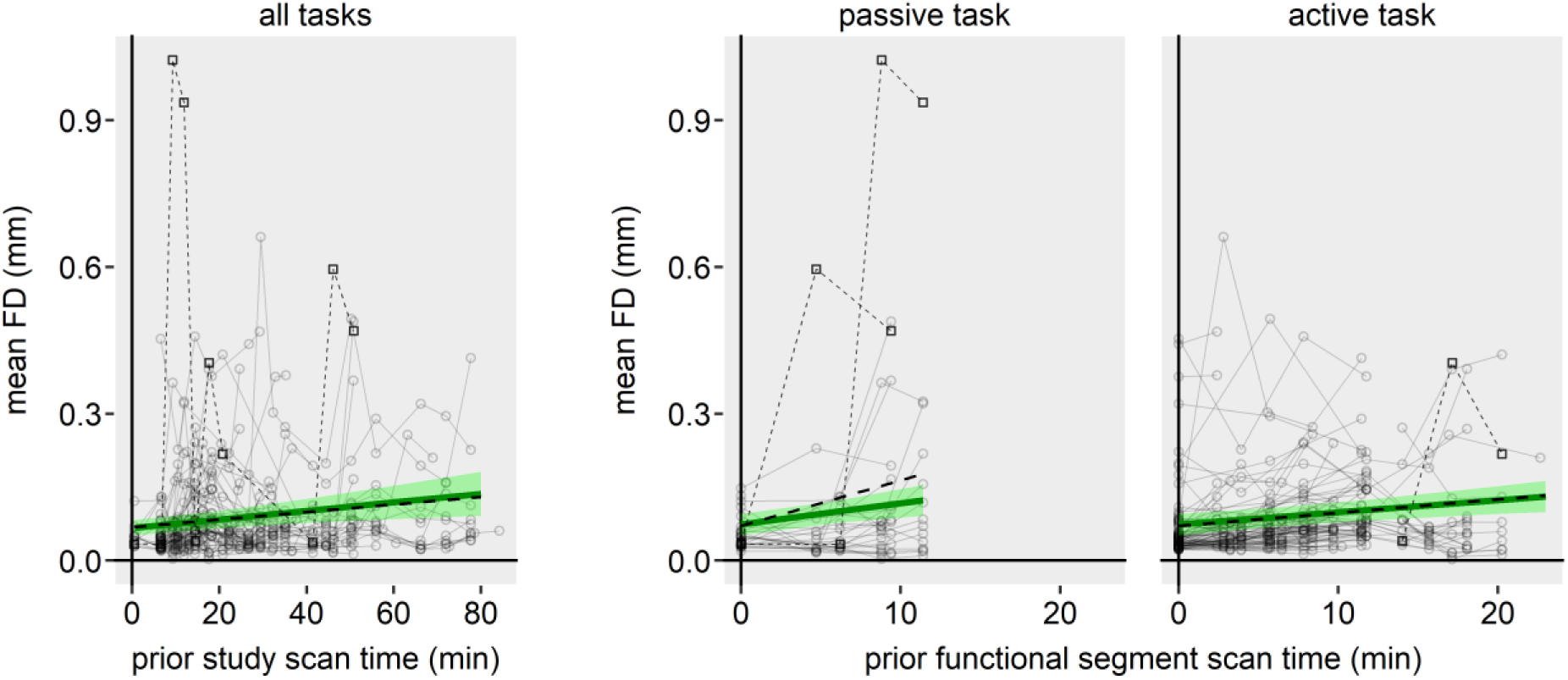
Predictor effect plots for adult’s head motion across the course of a study. Thick green lines are fitted linear regression lines for the fixed effects predictors denoted on the x-axes. Shaded green areas show the 95% confidence band for the regression lines. Each gray circle represents one run. Left: Head motion across the course of a study. Each circle-connecting line represents a participant‘s study. Right: Head motion across the course of a functional segment for passive and active fMRI tasks. Each circle-connecting line represents one participant’s functional segment data acquisition course. Participants that were scanned on multiple days, on multiple sessions, or with multiple functional segments are represented by multiple lines. Black dashed lines and squares show the participant that drove the task engagement × prior functional segment time interaction.

**Table 2.** Fixed effects parameter estimates of final model for adult’s mean motion across the course of a study.

| parameter | $\beta$ | Lower 95% CI | Upper 95% CI | SE | df | t | p |
| --- | --- | --- | --- | --- | --- | --- | --- |
| Intercept | 0.05030 | 0.03328 | 0.06732 | 0.00869 | 377 | 5.787 | < .001 |
| *Prior functional segment scan time* | 0.00250 | 0.00098 | 0.00401 | 0.00077 | 377 | 3.230 | 0.001 |
| *Prior study scan time* | 0.00087 | 0.00030 | 0.00144 | 0.00029 | 377 | 2.990 | 0.003 |
| *Task engagement* × *Prior functional segment scan time* | 0.00185 | -0.00058 | 0.00428 | 0.00124 | 377 | 1.491 | 0.137 |

Note: CI = confidence interval, SE = standard error, df = degrees of freedom

#### 3.1.3 Summary

Our results demonstrate that head motion in children and adults can be reduced by splitting up data acquisition. However, depending on the age group, different strategies seem to be effective. For children, our data suggests a benefit of splitting up data acquisition into multiple, short sessions on the same day and keeping the overall study length as short as possible. In contrast, adults benefited from interspersing experimental runs with inside-scanner breaks.

### 3.2 High-motion event occurrence increases with run length

#### 3.2.1 Children

*Minute of run* significantly predicted motion in children, i.e. children’s motion increased with increasing run length (Figure 7, Table 3). For example, in the third minute of a run, the average risk of a high-motion volume was at 16.5 %, while it increased to 18.3 % in the sixth minute of a run. *Prior day scan time* was also a significant predictor of motion. However, each value of this run-wise predictor affects each *minute of a run* equally and therefore does not contribute to explaining how motion develops over the course of a run. For example, *minute of run* 1, 2, and 3 always have the same values of *prior day time*. Thus, as the run-wise *prior day scan time* did not interact with minute of run, its effect is not of interest in this analysis.

**Figure 7:**
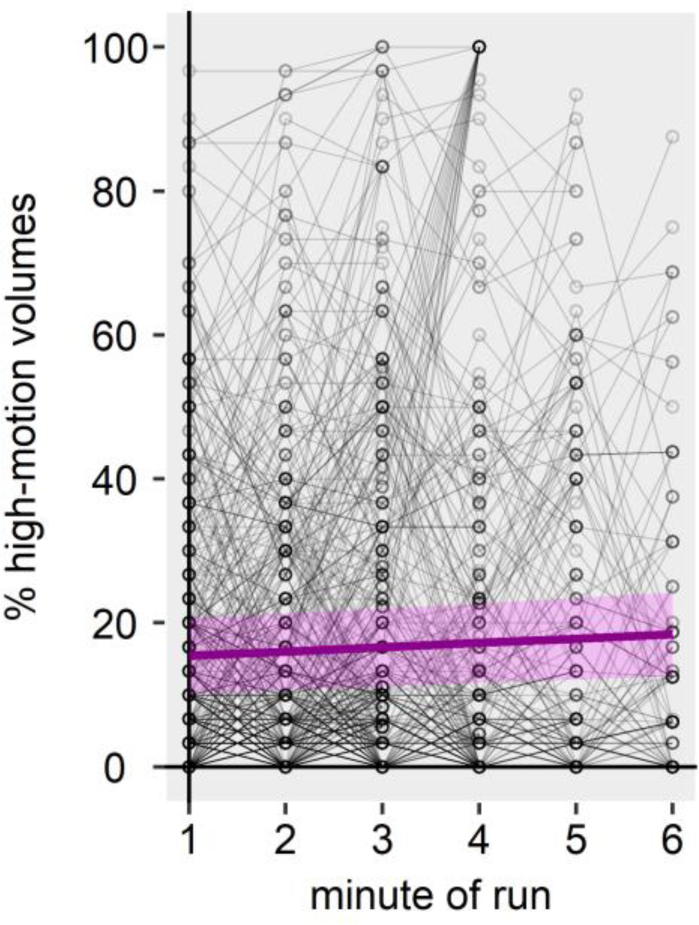
Children’s head motion across the course of a run. Each gray circle represents one minute bin. Each circle-connecting line represents a participant‘s run. Participants that were scanned for more than one functional run are represented by multiple lines. Thick magenta line shows the fitted linear regression line for the minute of run effect. Shaded magenta area show the 95% confidence band for the regression line.

**Table 3.** Fixed effects parameter estimates of final model for children’s frequency of motion peaks across the course of a run

| parameter | $\beta$ | Lower 95% CI | Upper 95% CI | SE | df | t | p |
| --- | --- | --- | --- | --- | --- | --- | --- |
| Intercept | 1.12857 | -2.87326 | 5.13041 | 2.04301 | 2290 | 0.552 | 0.581 |
| *Minute of run* | 0.60187 | 0.15491 | 1.04884 | 0.22818 | 2290 | 2.638 | 0.008 |
| *Prior functional segment scan time* | 0.22521 | -0.01436 | 0.46478 | 0.12231 | 2290 | 1.841 | 0.066 |
| *Prior session scan time* | 0.08717 | -0.18705 | 0.36138 | 0.13999 | 2290 | 0.623 | 0.534 |
| *Prior day scan time* | 0.41294 | 0.22250 | 0.60339 | 0.09722 | 2290 | 4.247 | < .001 |
| *Scanner training date* | 0.81829 | -0.71812 | 2.35470 | 0.78436 | 2290 | 1.043 | 0.297 |

Note: CI = confidence interval, SE = standard error, df = degrees of freedom

#### 3.2.2 Adults

*Minute of run* significantly predicted motion in adults, i.e. adult’s motion increased with increasing run length (Figure 8, Table 4). For example, in the third minute of a run, the average risk of a high-motion volume was at 5.3 %, while it increased to 6.5 % in the sixth minute of a run. *Prior functional segment scan time* and *prior session* scan *time* were also significant predictors of motion. However, as in children, these run-wise predictors did not contribute to explaining how motion develops over the course of a run as they did not interact with *minute of run* and thus are not of interest in this analysis.

**Figure 8:**
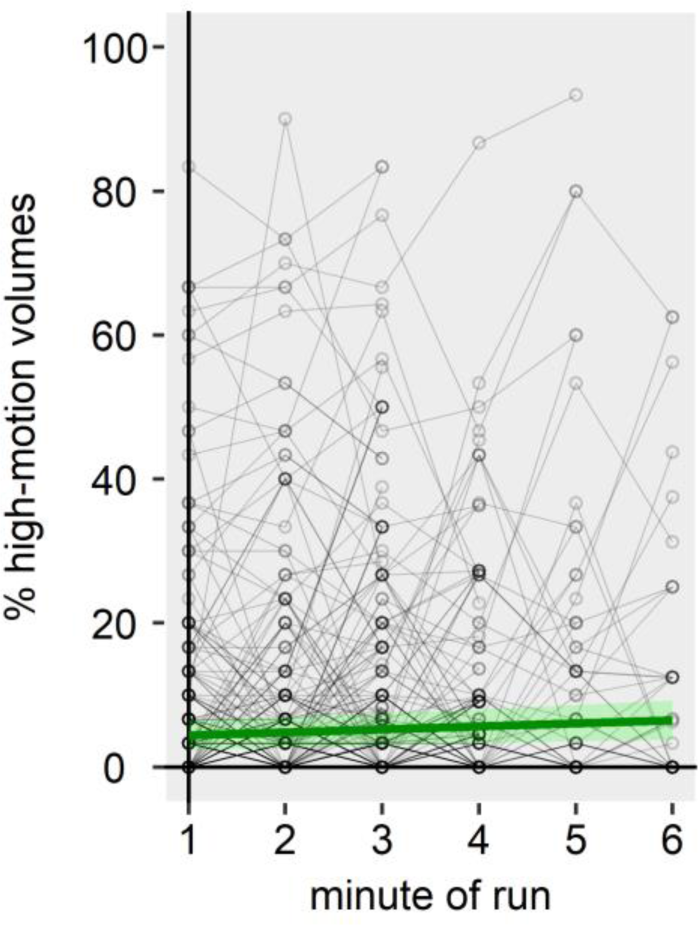
Adult’s head motion across the course of a run. Each gray circle represents a one minute bin. Each circle-connecting line represents a participant‘s run. Participants that were scanned for more than one functional run are represented by multiple lines. Thick green line shows the fitted linear regression line for the minute of run effect. Shaded green area show the 95% confidence band for the regression line.

**Table 4.** Fixed effects parameter estimates of final model for adult’s frequency of motion peaks across the course of a run

| parameter | $\beta$ | Lower 95% CI | Upper 95% CI | SE | df | t | p |
| --- | --- | --- | --- | --- | --- | --- | --- |
| Intercept | 0.80019 | -1.46836 | 3.06874 | 1.15779 | 1642 | 0.691 | 0.490 |
| *Minute of run* | 0.42562 | 0.10116 | 0.75008 | 0.16559 | 1642 | 2.570 | 0.010 |
| *Prior functional segment scan time* | 0.18341 | 0.00516 | 0.36165 | 0.09097 | 1642 | 2.016 | 0.044 |
| *Prior session scan time* | 0.11631 | 0.04504 | 0.18757 | 0.03637 | 1642 | 3.198 | 0.001 |

Note: CI = confidence interval, SE = standard error, df = degrees of freedom

#### 3.2.3 Summary

Our results indicate that long runs have a negative impact on data quality: The frequency of high-motion events increases with ongoing run length for both children and adults.

### 3.3 Control analysis: Age—but not motion—influenced the data acquisition procedure in children

Data acquisition in terms of which task was acquired when, the positions of structural scans, i.e. inside-scanner breaks, and the number of total runs was largely predetermined for each study and participant. However, we adapted our planning to the requirements of the given participant—mainly for children and only seldomly for adults. Specifically, participants were able to terminate the study, day or session at any moment. Most of the time, termination of a session was done after active inquiring about the well-being by the researchers. Here, children’s age might have influenced researchers’ sensitivity towards well-being reports and researchers’ decision to terminate. Crucially, subjective well-being (or boredom) might be associated with motion in the scanner. Thus, irrespective of reported well-being of the child, observed motion by the researchers (visible from the control room in Bangor and from standing next to the scanner bore in Bochum) might have influenced the decision to terminate the study, day, session, or functional segment early.

To investigate the possible effect of age and motion on the procedure, we ran separate multiple regression analyses for the total study length as well as the total number of acquired runs, sessions, and days using participants’ age and their mean percentage of high-motion-volumes as predictors. We chose the latter predictor over mean FD because only high-motion events would be visible for the researcher and act as a possible indicator to change the predefined study protocol.

The number of completed runs, sessions, or days was neither significantly predicted by age, nor by high-motion volumes (all *p*s for age > .0664; all *p*s for high-motion volumes > .1676). However, age—but not the mean percentage of high-motion volumes—was a significant predictor of total study length (age: β (SE) = 2.91 (1.29), *t* = 2.26, *p* = .027; high-motion volumes: β (SE) = −0.17 (0.16), *t* = −1.10, *p* = .275). Thus, it is unlikely that observable motion during fMRI scans was factored into the decision to deviate from our predefined study protocol, while young age might have been a factor.

## 4 Discussion

We identified factors that predict participant’s head motion in three neurodevelopmental fMRI studies including data of 77 children and 64 adults. Using MLMs, we investigated the effect of scanner training date, task engagement, as well as *prior scan time* since the beginning of the *study*, *day*, *session*, *or functional segment*. In children, splitting fMRI data acquisition into multiple sessions reduced motion. However, motion still increased across the course of a study, especially in older children. In adults, motion was reduced after task-free inside-scanner breaks but—as in children— motion still increased across the course of a study. In both children and adults, motion increased with run length.

### 4.1 Splitting up data acquisition reduces motion

#### 4.1.1 Children

Children’s head motion seems to increase across the course of a study, as *prior study scan time* predicted head motion, especially for older children. At the same time, we did not find that *prior day scan time* predicted head motion, suggesting that children do not seem to benefit from splitting up studies into several days. Alternatively, a possible effect of splitting up studies into several days was too small to be detected with the power of our study. To speculate on possible explanations, for children, the initial excitement of participating in an fMRI study and the commitment to do everything right might be less pronounced after the first day. On the second day, they might be less motivated to lie still, e.g. due to the lacking novelty of the situation (similar visual appearance or task demands).

Older children showed a larger increase in motion with increasing study length. Initially, this seems unexpected. Intuition would suggest that older children are better at lying still for extended periods of time. However, it seems that this *age* × *prior study scan time* interaction could be driven by a higher baseline motion of young children at the beginning of the study, which then does not change as much over time. In contrast, older children seem to be able to suppress the urge to move at the beginning of a study quite well, and then gradually relax into motion levels similar to that of younger children. Thus, we caution against overinterpreting this interaction as a cause to plan shorter studies just because older children are scanned, as predicted motion values at maximum study length were comparable and few studies will plan studies with more than 80 minutes of pure scan time.

Our results indicate that pediatric neuroimaging studies may benefit from breaking up data acquisition into several sessions on the same day, as children’s head motion increases with ongoing session length. So far, no other studies have investigated motion across multiple sessions on the same day. Nevertheless, single-session studies in children have found that motion increases with increasing session length in children (Achterberg and van der Meulen, 2019; Engelhardt et al., 2017). Our study points to an important distinction regarding the kind of breaks that researchers can schedule in children’s fMRI scanning protocols. While motion can be effectively reduced by allowing children to exit the scanner, inside-scanner breaks in fMRI data acquisition during which children cannot exit the scanner, but instead watch a video while a structural scan is acquired, do not seem to reduce motion after the break. Possibly, it is important for children to be able to move freely, get face-to-face social interactions, or just relieve the desire to void their bladder—none of which are possible without exiting the scanner. Alternatively, showing a video might have effects opposite to those desired: Because the video is highly engaging, it may lead to reduced motion during the break (Cantlon and Li, 2013; Vanderwal et al., 2015), but any subsequent normal fMRI task might be perceived as boring due to a contrast effect, leading to higher motion in turn.

In our experience, implementing outside-scanner breaks (i.e. splitting data acquisition into multiple sessions) is very feasible. In our three studies, breaks were designed to last 5 minutes outside of the scanner and we did not experience any problems with upholding this time limit. If a cover story for children is used, the break can be incorporated in that story (e.g. in a space journey theme there might be limited time for a space walk, maintenance, planet visit, etc.). Our 5-minute breaks usually prolonged the total study by 15 minutes due to getting participants out of and back inside the scanner, and running another survey scan. In contrast to longer breaks, such as 20 minutes or 2 hours, these short breaks limit the additional scanner costs and avoid both unfeasible study date arrangements with participants and parents as well as complicated scanner reservation arrangements. Thus, for studies with more than 30 minutes of total raw scan time, implementing a split into two sessions seems to be a feasible and effective way of reducing motion in children. Another approach is to schedule two or more children back to back, who then take turns in being in the scanner and taking breaks. It should be noted though that this approach might require additional staff to supervise children during scanning and breaks in parallel.

In our study, we did not find an effect of task engagement on head motion in children. This contradicts previous findings, which showed that children show less motion in engaging tasks in contrast to less engaging tasks or resting state (Engelhardt et al., 2017). This disparity might be due to different definitions and aims of the studies. In our analysis, we defined tasks as engaging if participants had to perform a task that included button-pressing, and we defined tasks as non-engaging if participants just had to passively watch the display. We chose this definition as our aim was to identify aspects in fMRI study designs that can be actively manipulated by the researcher. Engelhardt et al. (2017) defined engaging tasks as fast-paced, cognitively demanding, or socially engaging. These aspects are often inherent in a task and cannot be changed without changing the research question. For example, experiments on social development in children will need to have a socially engaging component. However, while researchers can decide if a given task will require a manual response or not, this choice alone does not seem to affect head motion. Possibly, motion-reducing effects due to engaging tasks that require button-pressing or high attention are also cancelled out by motion-increasing effects due to excitement or button-press-related motion.

The time between scanner training and the actual scan does not seem to be a major influence on children’s head motion during fMRI scans. Although scanner training timing improved (and thus was added to) the MLM, it did not emerge as a significant predictor in the final model. This assumption fits with the observed intraclass correlation for the grouping factor *study*. As all children in study C were trained on the same day as the actual fMRI scan but all children (except for one) in study A and B were trained between 1 and 28 (mean = 4.30) days before the actual fMRI scans, a high intraclass correlation for the grouping factor *study* would have indicated differences between studies (and thus sites) such as the scanner training procedure or the scanner training date. However, we observed an extremely low interclass correlation for the grouping factor *study*. Thus, planning scanner training as close to the actual scan as possible for strong recency effects, or planning scanner training some days before the actual scan to let it “sink in” seem to be equally effective (but also see 4.4, Future directions).

#### 4.1.2 Adults

For adults, we found that head motion seems to increase across the course of a study, as *prior study scan time* predicted head motion. While this result seems to mirror our findings in children, note that all but two adults were scanned on one day only—thus *prior study scan time* is equivalent to *prior day scan time* and our study cannot inform about possible benefits of splitting up data acquisition across days in adults. Interestingly, *prior functional segment scan time* predicted head motion in adults, while the *prior session scan time* did not contribute enough to the MLM to be included in the analysis. As motion across the day/study increased, this suggests that motion was reduced by inside-scanner breaks and that motion was seemingly constant across sessions. Consequently, the observed increasing motion across the course of a day is presumably due to a higher mean motion for the second session of the day. Thus, adults seem to be able to restrain their head motion across relatively long sessions, but are more likely to increase head motion in a second session on the same day. Therefore, in contrast to children, long studies in adults might best be acquired in one long session interspersed with inside-scanner breaks (possibly with short videos during anatomical scans). We are not aware of studies that investigated motion in adults across multiple sessions within the same day. However, our findings connect with the existing literature in so far as the number of completed runs in a single-session study with five runs did not have any effect on motion (Huijbers et al., 2017).

Regarding task engagement, we only found a very unstable interaction with *prior functional segment scan time* that was driven by on extreme outlier. Thus, it is doubtful that motion during passive tasks increased at a steeper rate with ongoing time after breaks. Moreover, we cannot complement previous findings that showed higher motion for passive tasks or resting state and lower motion for active tasks (Cantlon and Li, 2013; Huijbers et al., 2017; Vanderwal et al., 2015).

### 4.2 High-motion event occurrence increases with run length

We found that the frequency of motion peaks within a run increases with run length in both children and adults. This suggests that in general, it is preferable to plan rather short runs instead of longer runs. However, aside from statistically significance, our data also show that the increase per minute is moderate (0.60 % for children and 0.43 % for adults). Consequently, high-motion volumes do not just start to occur after a certain amount of time—they already occur in the first minute of the scan. In addition, most experimental paradigms will require a minimum duration of some sorts, e.g. to acquire a necessary number of volumes in order to have sufficient detection power or to have enough time for the hemodynamic response function to fully unfold a certain number of times (e.g. localizers), to achieve sufficient reliability (e.g. resting-state), or to map brain responses to stimuli of a certain length (movie segments). Thus, keeping runs as short as possible and acquiring high quality fMRI data should not compromise the quality of the experimental design. However, researchers are often free to choose if they plan to acquire the desired amount of data in a few long runs or in many short runs. For example, if 18 minutes of data should be acquired, instead of planning three 6-minute runs, we would suggest to plan five 3.6-minute runs. Based on our experience, while run durations around three minutes are still practicable in terms of scanner and experimental paradigm operation, we would not encourage run durations below 2 minutes.

Interestingly, we did not find interactions of *run minute* with any of the *prior scan time* variables. So, a common expectation—that motion peaks are especially evident at the beginning of the first run of a study or day—cannot be confirmed by our data. In a previous study that investigated the association between total run length and mean run motion in up to 6-7 minute long runs, results were mixed, i.e. a positive relationship in one sample and a negative relationship in the other sample (Engelhardt et al., 2017).

### 4.3 Limitations

While our study marks an important contribution towards understanding which factors are effective in obtaining good quality data from fMRI experiments, some limitations of our methodology should be considered. While the data acquisition procedure was predetermined for each study, participants were able to influence the *prior study time* factors and their head motion. Consider a child that shows increasingly observable motion and whose well-being reports indicate an increasing lack of interest after short periods of time. In consequence, we might have decided to implement more outside-scanner breaks than usual. This reaction would increase the number of sessions in this child and would also drive the effect of prior session scan time on motion.

However, our control analyses showed that observable motion during scans or children’s age did not influence the total number of completed runs, sessions, or days in children. Thus, while participants might have had influence on the exact timing of inside- or outside scanner breaks or the termination of a session, or day, the overall numbers of breaks was not biased. The only possible bias that our control analysis was able to reveal was that we might have been more attentive to the subjective well-being reports of younger children, more inclined to terminate young children’s studies’ early, or that younger children might have uttered the desire to terminate the study earlier, as age was a significant predictor of total study length.

With 626 runs from 77 children and 469 runs from 54 adults, our sample size is substantial. However, recent collaborative efforts such as the NCANDA, PING, or ABCD studies (Brown et al., 2015; Casey et al., 2018; Jernigan et al., 2016) have made larger pediatric neuroimaging datasets available. If detailed notes on inside- vs outside-scanner breaks are available in digital form, large open datasets could provide a powerful way of answering questions on how to prevent head motion. However, one downside of the highly standardized scanning protocols usually used in these large-scale multi-site studies is the lack of variability in scanning procedures (timing of breaks, number of runs, etc.) and the limited amount of fMRI data collected (if any non-resting-state data is collected at all).

### 4.4 Future directions

It remains unclear which contribution scanner training has on children’s head motion. Considering that a simple toy tunnel and a commercial mock scanner resulted in comparable success rates for high-quality structural images (Barnea-Goraly et al., 2014), it might be interesting to see if these results generalize onto strict motion thresholds during longer fMRI studies. Here, future studies should experimentally manipulate the kind of training for children, e.g. mock scanner training vs. toy tunnel training vs. no training. Also, these studies could investigate if short version trainings have comparable effects to extensive trainings sessions and at which age children cease to benefit from scanner training.

Aside from the investigated and discussed procedures, technical solutions might help to reduce motion. Recent developments have made real-time motion detection available that can be used to provide immediate feedback to participants and researchers (Dosenbach et al., 2017). In children younger than 10 years, providing immediate feedback has been successful in reducing motion using this technology (Greene et al., 2018). Thus, age effects in motion could occur not due to an inability to lie still in younger children, but possibly due to a lower awareness of their own movements. Even if motion is not reduced, this technology provides the possibility to adapt a common fMRI study protocol—based on group-average recommendations like those in this study—to the individual participant’s behavior.

This possibility might lead to a considerable reduction in cost and improvement of data quality, as our ICC analysis showed that motion differs substantially between *participants* and is relatively similar within participants. This finding agrees a previous studies finding that motion during fMRI tasks is a very stable neurobiological trait in children and adults that seems to be heritable, i.e. under strong genetic control (Achterberg and van der Meulen, 2019; Couvy-Duchesne et al., 2014; Engelhardt et al., 2017; van Dijk et al., 2012; Zeng et al., 2014).

### 4.5 Conclusion

The best way of dealing with head motion in fMRI is to prevent it in the first place. Our study shows that motion can be reduced by careful planning of the data acquisition procedure. Breaking up data acquisition into several sessions with the opportunity to leave the scanner is effective in reducing motion in children, while introducing inside-scanner breaks with continued acquisition of structural data is not effective. For adults, inside-scanner breaks during which further structural data can be acquired are a useful tool for preventing motion. Both children and adults benefit from short runs. To corroborate our findings of how study design can reduce and prevent motion in children and adults, future studies with an experimental approach are needed.

## Supporting information

Combined Supplementary Material

## Data and code availability statement

Data analysis code as well anonymized motion, demographic, and study procedure data is publicly available at the Open Science Framework (https://doi.org/10.17605/osf.io/jhm5t).

## Ethics statement

The of Ruhr University Bochum Faculty of Psychology and the University of Bangor School of Psychology ethics boards approved the studies, in which the data was acquired (Bochum: proposal no. 167 and 280, Bangor: proposal no. 2016-15743). All participants as well as children’s parents gave informed and written consent to participate voluntarily.

## Acknowledgements

We thank our teams of student assistants and interns for assisting during mock scanner training and data collection. We thank Katharina Limbach and Astrid Hönekopp for helpful discussions and comments on a previous version of this manuscript. We acknowledge the support of the Neuroimaging Centre of the Research Department of Neuroscience at Ruhr University Bochum and the Bangor Imaging Centre. We thank all participants and their parents for participating in our studies.

This work was supported by a PhD scholarship of the Konrad-Adenauer-Foundation and an International Realization Budget of the Ruhr University Bochum Research School PLUS through funds of the German Research Foundation’s Universities Excellence Initiative (GSC 98/3) to TWM, a Bangor School of Psychology funded PhD Scholarship to JW, a PhD scholarship of the German Academic Scholarship Foundation to MN, an ERC Starting Grant (Becoming Social, 716974) to KK, and grants from the Deutsche Forschungsgemeinschaft (DFG, German Research Foundation, project number WE 5802/1-1 and project number 316803389 – SFB 1280 project A16), the Mercator Research Center Ruhr (AN-2014-0056), and the Volkswagen Foundation (Lichtenberg Professorship, 97 079) to SW.

## Author contributions

Conceptualization: TWM; Methodology: TWM; Formal Analysis: TWM; Investigation: TWM, JW, MN; Resources: SW, KK; Data Curation: TWM, JW, MN; Writing—Original Draft: TWM; Writing—Review and Editing: TWM, MN, SW, JW, KK; Visualization: TWM; Supervision: SW, KK; Project Administration: TWM; Funding Acquisition: SW, TWM, JW, MN, KK

