## Supplementary material for "Head motion during fMRI tasks is reduced in children and adults if participants take breaks": Combined Supplementary Material

### Supplementary Figures

#### Figure S1

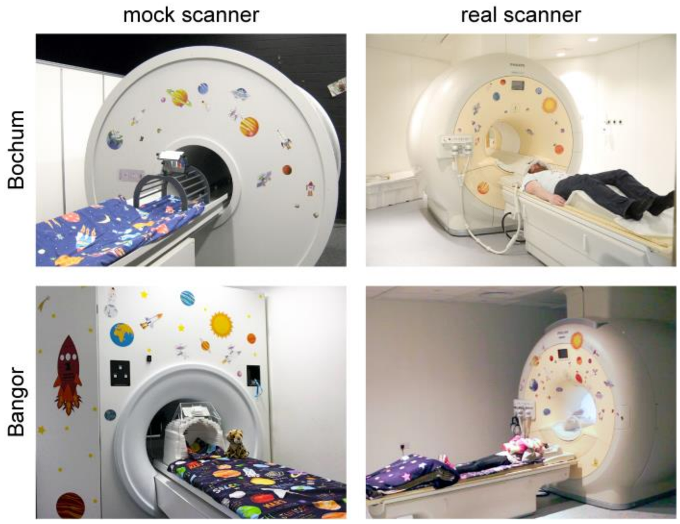

**Figure S1:** Mock scanner and real scanner at the two scanner sites in Bochum and Bangor.

### Supplementary Tables

#### Table S1

| task | participants | | runs | |
|---|---|---|---|---|
| | children | adults | children | adults |
| A1 | 33 | 15 | 132 | 60 |
| A2 | 27 | 15 | 108 | 60 |
| A3 | 16 | 14 | 46 | 42 |
| A4 | 26 | 13 | 74 | 38 |
| B1 | 12 | 25 | 18 | 50 |
| B2 | 17 | 25 | 60 | 96 |
| C1 | 27 | 14 | 81 | 42 |
| C2 | 27 | 14 | 80 | 40 |
| C3 | 27 | 14 | 27 | 14 |
| C4 | 0 | 9 | 0 | 27 |

#### Table S2

Table S2: Order of fixed effect parameter inclusion, given significant contribution to the mean FD model

| Test order | | Parameter |
|---|---|---|
| Child | Adult | |
| 1 | 1 | *Prior functional segment scan time* |
| 2 | 2 | *Prior session scan time* |
| 3 | - | *Prior day scan time* |
| 4 | 3 | *Prior study scan time* |
| 5 | - | *Age* |
| 6 | - | *Scanner training date* |
| 7 | 4 | *Task engagement* |
| 8 | - | *Age × Prior functional segment scan time* |
| 9 | - | *Age × Prior session scan time* |
| 10 | - | *Age × Prior day scan time* |
| 11 | - | *Age × Prior study scan time* |
| 12 | - | *Scanner training date × Prior functional segment scan time* |
| 13 | - | *Scanner training date × Prior session scan time* |
| 14 | - | *Scanner training date × Prior day scan time* |
| 15 | - | *Scanner training date × Prior study scan time* |
| 16 | 5 | *Task engagement × Prior functional segment scan time* |
| 17 | 6 | *Task engagement × Prior session scan time* |
| 18 | - | *Task engagement × Prior day scan time* |
| 19 | 7 | *Task engagement × Prior study scan time* |

### Table S3

Table S3: Order of fixed effect parameter inclusion, given significant contribution to the high-motion volume model

| Test order | | Parameter |
|---|---|---|
| Child | Adult | |
| 1 | 1 | *Minute of run* |
| 2 | 2 | *Prior functional segment scan time* |
| 3 | 3 | *Prior session scan time* |
| 4 | - | *Prior day scan time* |
| 5 | 4 | *Prior study scan time* |
| 6 | - | *Age* |
| 7 | - | *Scanner training date* |
| 8 | 5 | *Task engagement* |
| 9 | 6 | *Minute of run × Prior functional segment scan time* |
| 10 | 7 | *Minute of run × Prior session scan time* |
| 11 | - | *Minute of run × Prior day scan time* |
| 12 | 8 | *Minute of run × Prior study scan time* |
| 13 | - | *Minute of run × Age* |
| 14 | - | *Minute of run × Scanner training date* |
| 15 | 9 | *Minute of run × Task engagement* |

### Table S4

Table S4: Multilevel model creation process with significant additions to the simple intercept model for children's mean motion across the course of a study.

| No. | Model | df | AIC | LL | test | $\chi^2$ | *p* | $R^2$m | $R^2$c |
|---|---|---|---|---|---|---|---|---|---|
| 1 | Intercept | 2 | 110.14 | -53.07 | | | | | |
| 2 | Random intercept for *participant* and *session* | 4 | -258.71 | 133.35 | 1 vs 2 | 372.84 | < .001 | 0 | 0.692 |
| 3 | *Prior functional segment scan time* | 5 | -296.00 | 153.00 | 2 vs 3 | 39.29 | < .001 | 0.022 | 0.712 |
| 4 | *Prior session scan time* | 6 | -342.12 | 177.06 | 3 vs 4 | 48.12 | < .001 | 0.049 | 0.750 |
| 5 | *Prior day scan time* | 7 | -344.79 | 179.39 | 4 vs 5 | 4.67 | 0.031 | 0.056 | 0.758 |
| 6 | *Scanner training date* | 8 | -348.15 | 182.07 | 5 vs 6 | 5.36 | 0.021 | 0.067 | 0.766 |
| 7 | *Age × prior session scan time* | 9 | -351.36 | 184.68 | 6 vs 7 | 5.21 | 0.022 | 0.071 | 0.762 |
| 8 | *Age × prior study scan time* | 10 | -360.16 | 190.08 | 7 vs 8 | 10.80 | 0.001 | 0.084 | 0.775 |
| 9 | Random slopes for main effect predictors | 24 | -423.62 | 235.81 | 8 vs 9 | 91.47 | < .001 | 0.027 | 0.949 |

*Note*: df = degrees of freedom, AIC = Akaike's information criterion, LL = log(likelihood), $R^2$m = marginal $R^2$ (explained variance by fixed effects), $R^2$c = conditional $R^2$ (explained variance by fixed and random effects)

### Table S5

Table S5: Multilevel model creation process with significant additions to the simple intercept model for adult's mean motion across the course of a study.

| No. | Model | df | AIC | LL | test | $\chi^2$ | *p* | $R^2$m | $R^2$c |
|---|---|---|---|---|---|---|---|---|---|
| 1 | Intercept | 2 | -731.30 | 367.65 | | | | | |
| 2 | Random intercept for *participant* and *session* | 4 | -858.62 | 433.31 | 1 vs 2 | 131.32 | < .001 | 0 | 0.399 |
| 3 | *Prior functional segment scan time* | 5 | -876.35 | 443.18 | 2 vs 3 | 19.73 | < .001 | 0.028 | 0.436 |
| 4 | *Prior study scan time* | 6 | -882.11 | 447.05 | 3 vs 4 | 7.76 | 0.005 | 0.038 | 0.442 |
| 5 | *Task engagement × Prior functional segment scan time* | 7 | -895.84 | 454.92 | 4 vs 5 | 15.74 | < .001 | 0.067 | 0.470 |
| 6 | Random slopes for main effect predictors | 12 | -902.26 | 463.13 | 5 vs 6 | 16.41 | 0.006 | 0.068 | 0.530 |

*Note*: df = degrees of freedom, AIC = Akaike's information criterion, LL = log(likelihood), $R^2$m = marginal $R^2$ (explained variance by fixed effects), $R^2$c = conditional $R^2$ (explained variance by fixed and random effects)

### Table S6

Table S6: Multilevel model creation process with significant additions to the simple intercept model for children's frequency of motion peaks across the course of a run

| No. | Model | df | AIC | LL | test | $\chi^2$ | *p* | $R^2$m | $R^2$c |
|---|---|---|---|---|---|---|---|---|---|
| 1 | Intercept | 2 | 23813.61 | -11904.81 | | | | | |
| 2 | Random intercept for *participant*, *session*, and *run* | 5 | 22669.11 | -11329.56 | 1 vs 2 | 1150.50 | < .001 | 0 | 0.522 |
| 3 | *Minute of run* | 6 | 22664.06 | -11326.03 | 2 vs 3 | 7.06 | .008 | 0.001 | 0.523 |
| 4 | *Prior functional segment scan time* | 7 | 22561.03 | -11273.51 | 3 vs 4 | 105.03 | < .001 | 0.039 | 0.551 |
| 5 | *Prior session scan time* | 8 | 22468.00 | -11226.00 | 4 vs 5 | 95.03 | < .001 | 0.065 | 0.556 |
| 6 | *Prior day scan time* | 9 | 22461.78 | -11221.89 | 5 vs 6 | 8.22 | 0.004 | 0.084 | 0.567 |
| 7 | *Scanner training date* | 10 | 22444.32 | -11212.16 | 6 vs 7 | 19.46 | < .001 | 0.101 | 0.586 |
| 8 | Random slopes for main effect predictors | 30 | 22283.94 | -11111.97 | 7 vs 8 | 200.38 | < .001 | 0.039 | 0.879 |

*Note*: df = degrees of freedom, AIC = Akaike's information criterion, LL = log(likelihood), $R^2$m = marginal $R^2$ (explained variance by fixed effects), $R^2$c = conditional $R^2$ (explained variance by fixed and random effects)

### Table S7

Table S7: Multilevel model creation process with significant additions to the simple intercept model for adult's frequency of motion peaks across the course of a run

| No. | Model | df | AIC | LL | test | $\chi^2$ | *p* | $R^2$m | $R^2$c |
|---|---|---|---|---|---|---|---|---|---|
| 1 | Intercept | 2 | 15093.84 | -7544.92 | | | | | |
| 2 | Random intercept for *participant*, *session*, and *run* | 5 | 13909.88 | -6949.94 | 1 vs 2 | 1189.96 | < .001 | 0 | 0.645 |
| 3 | *Minute of run* | 6 | 13907.03 | -6947.51 | 2 vs 3 | 4.85 | 0.028 | 0.001 | 0.646 |
| 4 | *Prior functional segment scan time* | 7 | 13872.96 | -6929.48 | 3 vs 4 | 36.07 | < .001 | 0.017 | 0.649 |
| 5 | *Prior session scan time* | 8 | 13861.00 | -6922.50 | 4 vs 5 | 13.96 | < .001 | 0.02 | 0.650 |
| 6 | Random slopes for main effect predictors | 17 | 13749.38 | -6857.69 | 5 vs 6 | 129.62 | < .001 | 0.02 | 0.699 |

*Note*: df = degrees of freedom, AIC = Akaike's information criterion, LL = log(likelihood), $R^2$m = marginal $R^2$ (explained variance by fixed effects), $R^2$c = conditional $R^2$ (explained variance by fixed and random effects)

### Table S8

Table S8. Fixed effects parameter estimates of final model for adult's mean motion across the course of a study before the exclusion of one outlier participant.

| parameter | β | Lower 95% CI | Upper 95% CI | SE | df | t | *p* |
|---|---|---|---|---|---|---|---|
| Intercept | 0.05005 | 0.03062 | 0.06948 | 0.00992 | 385 | 5.043 | < .001 |
| *Prior functional segment scan time* | 0.00271 | 0.00107 | 0.00435 | 0.00084 | 385 | 3.244 | 0.001 |
| *Prior study scan time* | 0.00078 | 0.00023 | 0.00132 | 0.00028 | 385 | 2.787 | 0.006 |
| *Task engagement × prior functional segment scan time* | 0.00662 | 0.00347 | 0.00978 | 0.00161 | 385 | 4.110 | < .001 |

*Note*: CI = confidence interval, SE = standard error, df = degrees of freedom

### Supplementary Text

#### Text S1

Study A included four tasks. Task A1 presented photographs of scenes, objects as well as gray rectangles at a rate of 1 Hz for a total of up to four 3.23-minute runs. The three categories were presented in a block-wise fashion for a total of thirteen 14-second blocks. Participants performed a 1-back task, i.e. they had to press a button whenever an image was presented twice in a row. For detailed information on Tasks A1, see Meissner et al. (2019). Task A2 presented photographs of familiar and unfamiliar scenes, as well as gray rectangles, at a rate of 0.5 Hz for a total of up to four 2.80-minute runs. The three categories were presented in a block-wise fashion for a total of thirteen 12-second blocks. Participants were instructed to press a button, whenever a small green fly was present in an image. Task A3 and Task A4 presented photographs of houses for a total of up to three 5.73-minute runs. Images were presented in pairs, but subsequently, i.e. an image was presented for 800 ms, followed by an inter-stimulus-interval of 400 ms, followed by the second image for 800 ms. The second image was the same image as the first, a different version of the first image, or a different image. The next image would be presented after a jittered inter-trial-interval of 2000-4000 ms. Again, participants were instructed to press a button, whenever a small green fly was present in an image.

Study B included two tasks. Task B1 presented photographs of faces, objects as well as unicolor rectangles at a rate of 0.5 Hz for a total of up to two 2.40-minute runs. The three categories were presented in a block-wise fashion in a total of eleven 12-second blocks. Participants were instructed to press a button whenever an image was blue-washed. Task B2 presented faces and scrambled faces at a rate of 0.5 Hz for a total of up to four 3.92-minute runs. The two categories were presented in a block-wise fashion in a total of seventeen 12-second blocks. Face-blocks could be one of three conditions. 1) A single image of one person’s face was presented repeatedly throughout the block. 2) Different images of the same person’s face were presented throughout the block. 3) Different faces of different people were presented throughout the block. Again, participants were instructed to press a button whenever an image was blue-washed. For detailed information on Tasks B1 and B2, see Nordt et al. (2018).

Study C included four tasks. Task C1 consisted of brief videos of point-light figures interacting, not interacting, or scrambled versions of interacting figures, presented in 16- second blocks for a total of three 2.60-minute runs. Task C2 consisted of brief videos of moving faces, bodies, and objects (see Pitcher et al., 2011), presented in 18- second blocks for a total of three 4.70-minute runs. Task C3 consisted of a single 6.20-minute animated video that has been previously used to identify mentalizing responses by contrasting time-points that evoke spontaneous mentalizing, and contrasting them with control time-points (see Richardson et al., 2018). For tasks C1-C3, participants were instructed to passively view stimuli without making any button-press responses; see Walbrin et al. (2020) for detailed information on these tasks. Finally, task C4 consisted of brief videos of point-light figures performing everyday biological movements (e.g. jumping), rotating point-light objects, and scrambled versions of these stimuli presented in 18s blocks for a total of three 3.13-minute runs, while subjects performed a 1-back task.

#### Text S2

MLMs were created in a data-driven process. First, we assessed the possibility that the grouping variables would introduce dependencies in the data—and thus confirm the need for an MLM. To this end, we calculated the intraclass correlation (ICC) for each grouping level of the model. The ICC is the proportion of the total variance that is explained by the respective grouping factor—in other words, the correlation between two randomly selected observations from within the same grouping factor, for example $ICC_{study} = \sigma^2_{study} / (\sigma^2_{study} + \sigma^2_{participant} + \sigma^2_{day} + \sigma^2_{session})$, where $\sigma^2_{grouping\ level}$ denotes the variance within the respective grouping level. A high ICC points to a relevant grouping factor, while grouping factors with low ICC can be ignored as they do not have any influence on the data. That is, observations within these grouping factors are not more similar to each other than observations between these grouping factors. We incorporated all grouping levels into our model that would explain at least 1% of the total variance. This criterion was determined after ICC calculations and upon inspection of the ICC distribution for all levels. Next, we tested if the model that incorporated the grouping structure actually had a better fit to our data.

#### Text S3

We used maximum-likelihood estimation to fit a linear baseline model to the data that predicted FD using the intercept, i.e. the mean FD, only. Then, we fit the same model, but allowed the intercept to vary (random intercepts) over all included grouping levels (e.g. different intercepts for each participant and session) and tested if this new model would be a better fit for our data. 1) The change in the -2×log(likelihood) (-2LL) between the models, determined by a chi-square likelihood ratio test, had to be significant at a threshold of $\alpha = .05$. 2) The Akaike's information criterion (punishing the -2LL for model complexity; $AIC = -2LL + 2 \times n_{\text{parameters in the model}}$) for the model that incorporated the grouping structure also had to indicate a better model fit, i.e. be smaller than for the previous model. If the two criterions were met, random intercepts for the grouping structure were included in all subsequent models, otherwise, the grouping structure was ignored.
